# Homeostatic Control on the Thought: a Comprehensive Explanation of Mind Wandering

**DOI:** 10.1101/2024.04.19.590376

**Authors:** Kazushi Shinagawa, Kota Yamada

**Affiliations:** Department of Psychology, Keio University, Tokyo, JAPAN; Keio University Global Research Institute, Tokyo, Japan; Department of Information Medicine, National Institute of Neuroscience, National Center of Neurology and Psychiatry, Japan; Institute for Quantitative Biosciences, The University of Tokyo, Tokyo, JAPAN

## Abstract

Our thoughts are inherently dynamic, often wandering far from the current situation. Mind-wandering (MW), which is these thought transitions, is crucial for understanding the nature of human thought. Although previous research has identified various factors influencing MW, a comprehensive framework integrating these findings remains absent. Here, we propose that homeostasis has the potential to explain MW and validate the idea through simulations by replicating previous findings. We employed a homeostatic reinforcement learning model where independent drives for the task and others were assigned, and drive reduction became a reward and trained under sustained attention to the response task. We confirmed that the model behaves consistently with the empirical results reported in human experiments, suggesting that the model accurately captures the underlying mechanism of MW. Finally, we discuss the behavioral and neurobiological commonality between human thought and animal behavior and the possibility that the same principle, homeostasis, controls these phenomena.

## Introduction

The thoughts of human beings are highly dynamic, and we often think about what is disengaging from the current situation, such as events that occurred in the past, might happen in the future, or will never happen. These thoughts shift from the primary task to unrelated thoughts, termed mind-wandering (MW; Smallwood and Schooler, 2015). Since MW is an ordinary phenomenon in which we spend 30∼50% of our waking time, it has important roles in human cognitive function (Christoff et al., 2016; Killingsworth and Gilbert, 2010).

Various factors encourage or discourage us to engage in MW. The first is the participants’ motivation towards the main task and task-unrelated thoughts (TUTs). As the task proceeds, participants’ motivation declines, leading to more MW and poorer task performance (Brosowsky et al., 2020; Zanesco et al., 2024). Motivation for internal thought also influences MW. The content of TUT often directs to the future, suggesting that individual worries and concerns drive MW (Baird et al., 2011; Poerio et al., 2013; Smallwood and Schooler, 2006). The task difficulty has a non-monotonic effect on MW. When the task is easy, it allows participants to engage in TUT and as the difficulty increases, engagement in TUT decreases (Seli et al., 2018; Thomson et al., 2013). Conversery, when the task is too difficulty, participants turned to engage in TUT more (Barrington and Miller, 2023; Kahmann et al., 2022; Xu and Metcalfe, 2016). Although various factors are involved in MW, and several hypotheses have been proposed to explain them, a comprehensive framework that unifies these findings is still absent.

Once we look at the daily life of animals, including humans, we can see that organisms engage in various behaviors. It is not limited to daily lives, and they come in experimental situations in both animals and humans (Breland and Breland, 1961; Eisenberger et al., 1967; Falk, 1966; Gentry, 1968; Kachanoff et al., 1973; Levitsky and Collier, 1968; Skinner, 1948). These facts remind us that there are similarities between thoughts and behaviors. Thus, we can assume that a common basis implementing thought transition, which seems unique to humans, and the behavior of organisms. A primordial source that drives animals to engage in specific behavior is physiological requirements, such as hunger and thirst. The idea of maintaining a steady physiological state, or homeostasis, has been applied to associative learning, habituation and sensitization, social behavior, and various behavioral phenomena (Eisenstein and Eisenstein, 2006; Juechems and Summerfield, 2019; Keramati and Gutkin, 2014; Lee et al., 2021; Uchida et al., 2022), suggesting that it has a potential to be applicable to human thoughts.

Homeostatic reinforcement learning incorporates the idea of homeostasis in how agents make decisions to keep their physiological state stable(Keramati and Gutkin, 2014). This model defines reward as the proximity to a desired value for the internal state (i.e., setpoint). Therefore, the reward changes dynamically according to the physiological state of the agent. For example, when an animal is hungry, food reduces its hunger and reinforces its behaviors. However, if the animal is full, excessive foods make the animal nausea and punish its behavior. HRL is a model learning the policy to prevent the physiological state from diverging by modeling changes in the reward according to the setpoint.

In this study, we replicated earlier findings by applying sustained attention to response task (SART), widely used in MW research, to HRL agents. In Simulation 1, we showed that MW occurs in HRL agents during the task and analyzed the model’s internal variables to identify the mechanism underlying MW occurrence in the model. Subsequently, we qualitatively replicated results of existing studies to validate use of HRL as a comprehensive framework for MW. In Simulation 2, we manipulated the parameter that determined the motivation for the main task and replicated that changes in motivation affect the proportion of MW. In Simulation 3, we replicated results that highly motivated events drive TUT by manipulating the motivation for TUT. In Simulation 4, we manipulated the task difficulty and replicated the findings that MW decreases as the task becomes more difficult but increases when the difficulty reaches an extreme level. Additionally, we discussed that these results imply that human-specific phenomena, like thought dynamics, are also governed by biological principles across physiology and behavior.

## Results

In the present study, we attempted to replicate the results reported in the MW study by conducting SART on agents implementing HRL (Keramati and Gutkin, 2014). In SART, fixation and numbers are presented alternately, and participants are required to respond only when a specific number is presented (Fig. 1A). It has been reported that response time to the target stimulus and its variance is increased during MW (Henríquez et al., 2016; Irrmischer et al., 2018; Leszczynski et al., 2017; Makovac et al., 2019; Seli et al., 2013; Shinagawa et al., 2023; Smallwood et al., 2008). We simplified this task to include only “fixation” and “stimulus” presentations for applying this task to agents (Fig. 1B). HRL agents could choose an action at each time step from three options: “response,” “focus,” and “task-unrelated thoughts” (TUT). The “response” indicates a response to the stimulus, the “focus” indicates that the agent is engaged in the task but does not respond, and “TUT” indicates that the agent is engaged in the TUT. In this simulation, we defined MW as transitioning from a task-focused state to TUT and its maintenance during the task. We constrained the agents’ choice depending on the environment state and previous action so that agents would not choose “response” when the agents engaged in “TUT” or the environment state is “fixation” (Fig 1C; See Materials and Methods for detail). In the simulations, one trial involved 40 timesteps that were composed of 30 “fixation” steps followed by 10 “stimulus” steps, and each agent conducted 200 trials. The homeostatic state of “TUT” is equal to the setpoint from the onset of the task, meaning that the drive for “TUT” always becomes negative during the task. Agents choose an action at each time step based on the past rewards and actions and calculate the reward and reward prediction error (RPE) based on the observations and their homeostatic state (Fig 1D; See Materials and Methods for detail). The following four simulations share the same environment, model architecture, interaction between agent and environment, and the flow of the simulation.

**Fig. 2.**
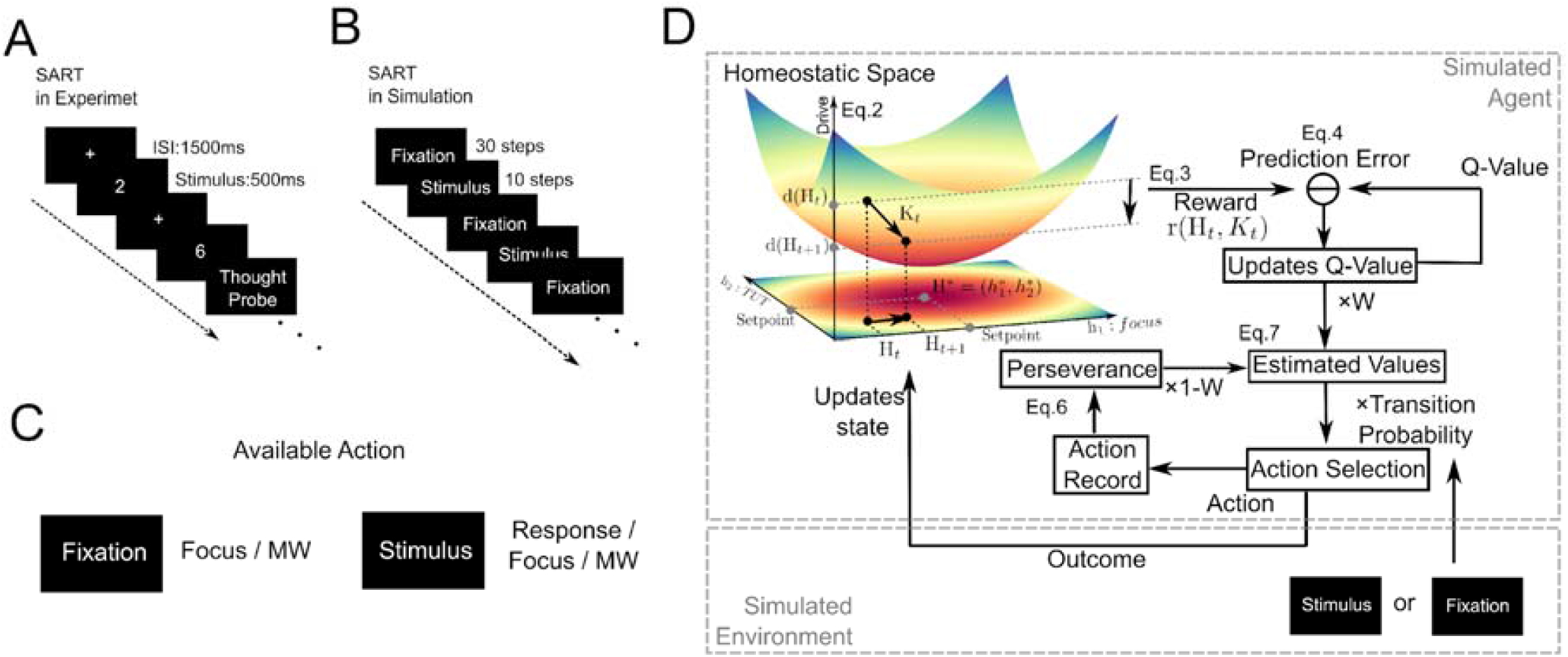
The task and agent used in the simulation. A. Schematic image of a typical SART recruited in an empirical MW study. B. The image of SART in the simulation. The duration of the “stimulus” presentation was set to be similar to the actual task conditions, with each step set to be 50 ms. C. The actions that would be selected under each task state in the simulation: During “fixation,” the agents could not respond and must either focus on the task and wait or be immersed in TUT. The agents can respond during “stimulus” presentation only if focused on the task. See Materials and Methods for details. D. Schematic image of HRL with a two-dimensional homeostasis space: When agents choose an action, agents receive an observation from the environment. The homeostatic state changes based on the observation and the deviation from the setpoints are calculated. This proximity is defined as a reward. The model calculates RPE by comparing the reward and the expected reward obtained by the chosen action, Q-value. RPE signal then updates the Q-value for the action. See Materials and Methods for details.

### Simulation1:Exploration of the process by which MW occurs

The goal of simulation 1 is to show that the occurrence of MW depends on two independent processes: 1) a choice between “focus” and “TUT,” and 2) independent dynamics of the drive are behind MW. To achieve this goal, we simulated and analyzed the behavior of three models: a full model incorporating both processes, a no-TUT model, and a vanilla Q-learning model (VQ). The no-TUT and VQ are models that lacked either of two processes from the full model (See Materials and Methods for details). Comparing the proportion of each action, “TUT” was rarely chosen other than the full model (Fig 2A). Response times in trials when agents engaged in “TUT” immediately before the “stimulus” onsets were prolonged in the full model and VQ than in trials when agents engaged in “focus” (Fig 2B). Furthermore, as pointed out in many recent studies (Leszczynski et al., 2017; Seli et al., 2013; Shinagawa et al., 2023), the variance of response time was larger during “TUT” than the “focus” only in the full model (Fig 2C). In addition, the VQ’s response times were scattered around 50 ms when the agents selected “TUT” upon presenting a “stimulus” (Fig 2B). Considering that each time step is 50 ms and that the agents cannot choose “response” at the “fixation” (Fig 1C), the agents chose “response” at 50 ms is the fastest way from “TUT.” In the VQ agents, the probability of “TUT’’ engagement rapidly decreased from the start of the session and rarely occurred from the middle phase of the session (Fig. 2D). These results suggest that two independent processes, 1) a choice between “focus” and “TUT,” and 2) independent dynamics of the drive behind them, are necessary to replicate the pattern of response times and shifts in thought observed in typical SART.

**Fig. 2.**
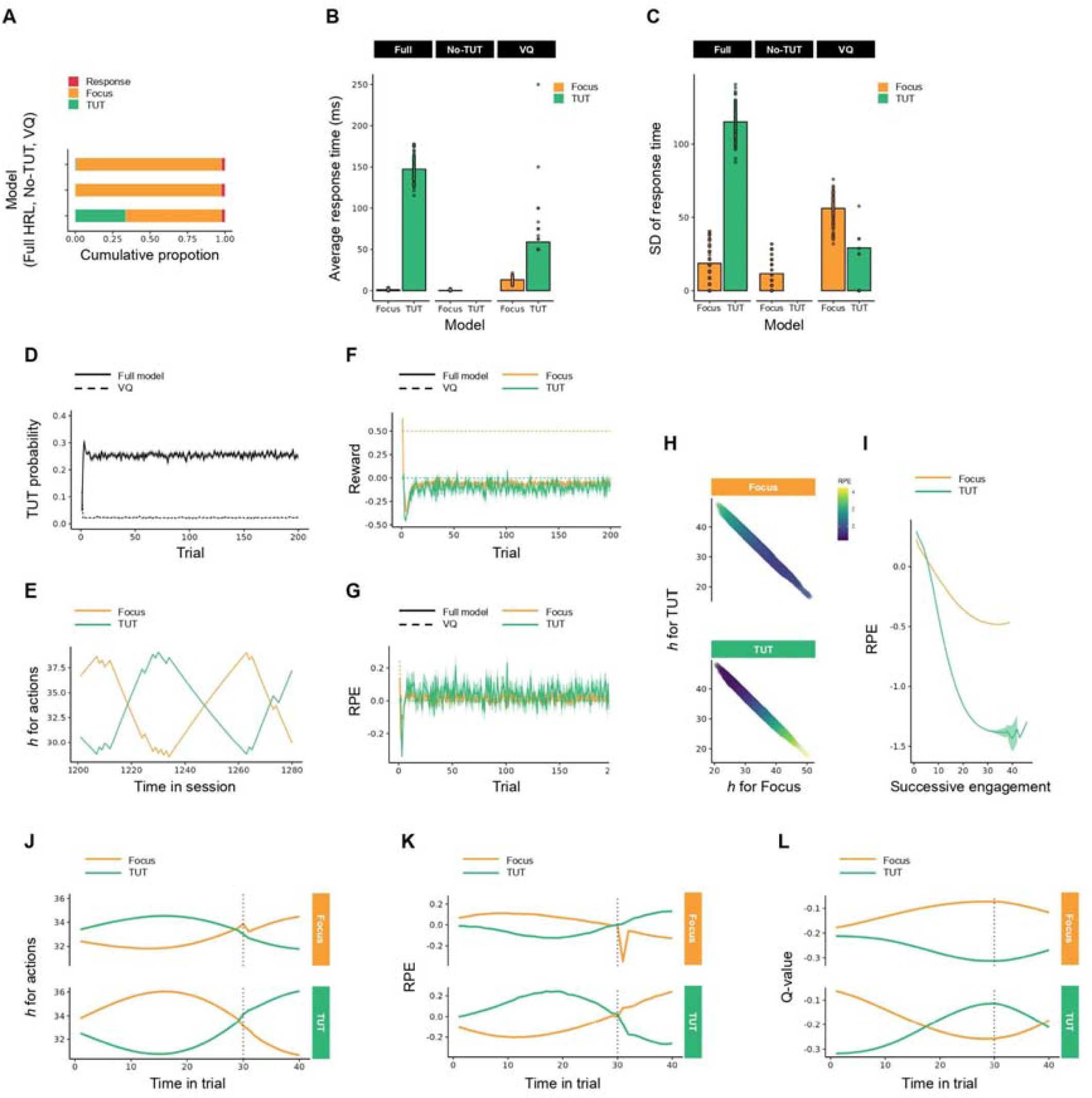
Results of simulation 1 A. Proportion of three actions, “response” (red), “Focus” (orange), and “TUT” (green), simulated by three models, the full model (bottom), no-TUT model (middle), and Vanilla Q-learning (top). B. Bar plots show response time from “stimulus” onset, averaged across agents, for each response immediately before “stimulus” onset. Because agents could not take “response” in the “fixation” period, we showed results that agents took “focus” or “TUT” immediately before “stimulus” onset. Individual panels show the results from each of the three models (from left to right: full model, no-TUT, VQ). Scattered dots above and below the bars indicate results from individual agents. C. Bar plots show standard deviation of response time immediately before “stimulus” onset among agents for each three models. D. TUT probability, averaged across agents, at each trial throughout the session for the full model (solid line) and VQ (dashed line). The grayed shades indicate standard error across agents. E. An example of the dynamics of the homeostatic state, *h*, for “focus” (orange) and “TUT” (green) in the full model. The data from 1200 timesteps to 1280 timesteps for one agent is shown. F. Trial-by-trial changes in rewards obtained by “focus” (orange) and “TUT” (green) simulated by the full model (solid line) and VQ (dashed line). G. Trial-by-trial changes in RPEs occurred by “focus” (orange) and “TUT” (green) simulated by the full model (solid line) and VQ (dashed line). H. The relationship between homeostatic states, *h*, for “focus” and “TUT” and RPEs generated by each two actions. The scatter plots show the RPEs resulting from the chosen action in the homeostatic state of the “focus” and “TUT” at a given time step. The upper panel and bottom panel show the RPEs generated by “focus” and “TUT” respectively. As the point color becomes brighter, the magnitude of RPE increases. I.The lines show that changes in RPEs are caused by successive engagement in the same action. We calculated the number of times “focus” or “TUT” is chosen, and averaged RPEs for each time across agents. J–L. Within-trial dynamics of homeostatic state, *h* (J), RPEs (K), and q-values (L), for “focus” (orange) and “TUT” (green) averaged over agents and trials. The upper and bottom panels show the data from the trial where agents choose “focus” and “TUT” immediately before stimulus onset, respectively.

We analyzed the model’s internal variables, such as homeostatic states, q-values, and RPEs, during the task to elucidate the mechanism by which MW occurs only in the full model. In HRL, rewards and RPEs for “focus” and “TUT” varied throughout the session, while they remained constant except during the initial period in VQ (Fig 2F, G). RPEs systematically changed depending on the homeostatic state of each action and the other (Fig. 2E, H). Specifically, the RPEs for the “focus” increased with the homeostatic state of “focus” decreased, and that for the “TUT” increased (Fig. 2H top). This relationship was reversed for “TUT” (Fig. 2H bottom). Furthermore, we analyzed changes in RPEs through successive engagement for a specific action. RPEs were initially positive at the onset of engagement for “focus” and “TUT,” but they decreased as successive engagement in each action and turned to be negative (Fig 2I). Successive engagement in a specific action led to saturation for the action and punished it. In contrast, unsaturated actions reduced the saturation of different actions and acquired a rewarding effect. In trials in which the agents were engaged in the “TUT” upon “stimulus” onset, the saturation in the homeostatic state of “focus” during the “fixation” period caused the reward and RPE to turn negative, which reduced the q-value (Fig. 2J, K, L). The saturation of the homeostatic state of the “focus” was alleviated by engaging in “TUT,” which turned the RPEs of “TUT” positive and increased the q-value. These results indicated that MW was caused by punishment due to saturation of “focus” and rewarding “TUT” due to reduced saturation, and these relationships were reversed in the “TUT” trials. In summary, MW continuously occurs throughout the session in HRL, as “focus” and “TUT” alternated to reduce the divergence of homeostatic states for each behavior (Fig. 2E).

In simulation 1, comparing the HRL including “TUT” as an option of action with two models that omit specific processes revealed that the increased response time and its variance and the proportion of “TUT” engagement observed in the SART stems from the choice between task-related actions and “TUT” engagement and distinct dynamics of the drive behind them. Examining the model’s internal variables revealed that MW is driven by two processes: the homeostatic state becomes saturated due to successive engagement in “focus,” and this saturation is alleviated through engagement in “TUT.” The same process also mediates the change in choice from “focus” to engaging in “TUT,” indicating that HRL replicated the cycle of attention during SART.

### Simulation2∼4: Qualitative replication of MW study using HRL

To demonstrate that HRL could unify previous findings and provide a comprehensive explanation for MW, we performed the following four simulations in which we manipulated the parameters of HRL to replicate the results reported in empirical studies qualitatively. In Simulations 2-1 and 2-2, we replicated the results that subjective motivation to the task affects the behavioral features, such as the “TUT” proportion, response time, its variance, “focus” and “TUT” sustainabilities, and their occurrence frequency by manipulating the setpoint (2-1) and the time decay of the homeostatic state (2-2) for the task-related actions. In Simulation 3, we replicated the hypothesis that highly interested concerns trigger MW (Baird et al., 2011; Poerio et al., 2013; Smallwood and Schooler, 2006). This was achieved by adjusting the setpoint for “TUT” to reflect varying interest in the MW content. In Simulation 4, we replicated the finding that MW was less likely to occur when task difficulty was high and more likely to occur when the task difficulty was extremely high (Barrington and Miller, 2023; Kahmann et al., 2022; Seli et al., 2018; Thomson et al., 2013; Xu and Metcalfe, 2016). Assuming that the reward presentation probability corresponds to the task difficulty, we examined the effect of manipulating the difficulty on the MW in the HRL model. Except for the manipulated parameters, the experimental environment and model parameters were the same as in Simulation 1.

In Simulation 2-1, to replicate the findings that MW is more likely to occur with decreasing motivation for the task, we simulated the behavior when the setpoints for the task-related actions were manipulated. First, we revealed that the HRL agents also engaged more “TUT” with decreasing the motivations for the task-related actions (Fig. 3A). While the response time and variance were not changed with motivation level when the agents engaged in “focus” upon the “stimulus” presentation, these were increased when the agents engaged in “TUT” (Fig. 3B, C). When actions occur as clusters, called bouts, there are different aspects of behavior, such as interbout interval and number of actions involved in each bout. To clarify how setpoint manipulation influenced which aspects of behavior, we analyzed the microstructure of behavior. We counted the number of repetitions from the onset of action to the end and revealed that motivations for task-related actions did not affect the average successive engagement for “focus” but decreased “TUT” (Fig 3D). On the other hand, the average interbout interval, intervals between the ends of action to the next onsets, decreased in “focus” but increased in “TUT” with increasing motivation levels (Fig. 3E). These results suggest that the higher setpoint shortened the successive engagement for “TUT” and it made response time shorter. To clarify how setpoint manipulation influenced the successive engagement of “TUT”, we conducted further analysis of the internal variables of the agents. When the setpoint was higher, the homeostatic state converged at a higher level, leading to enhanced attenuation (Fig. 3F, G) due to its dependence on the current homeostatic state and time decay (Eq. 1). This relevancy between setpoint and time decay suggests that recovery from saturation was faster in the higher setpoint condition. When the homeostatic state for “focus” was saturated, agents shifted to engage in “TUT” until the state of “focus” recovered from saturation. Thus, when the setpoint was higher, the homeostatic state for “focus” could recover from saturation faster, resulting in the short successive engagement in “TUT.”

**Fig. 3.**
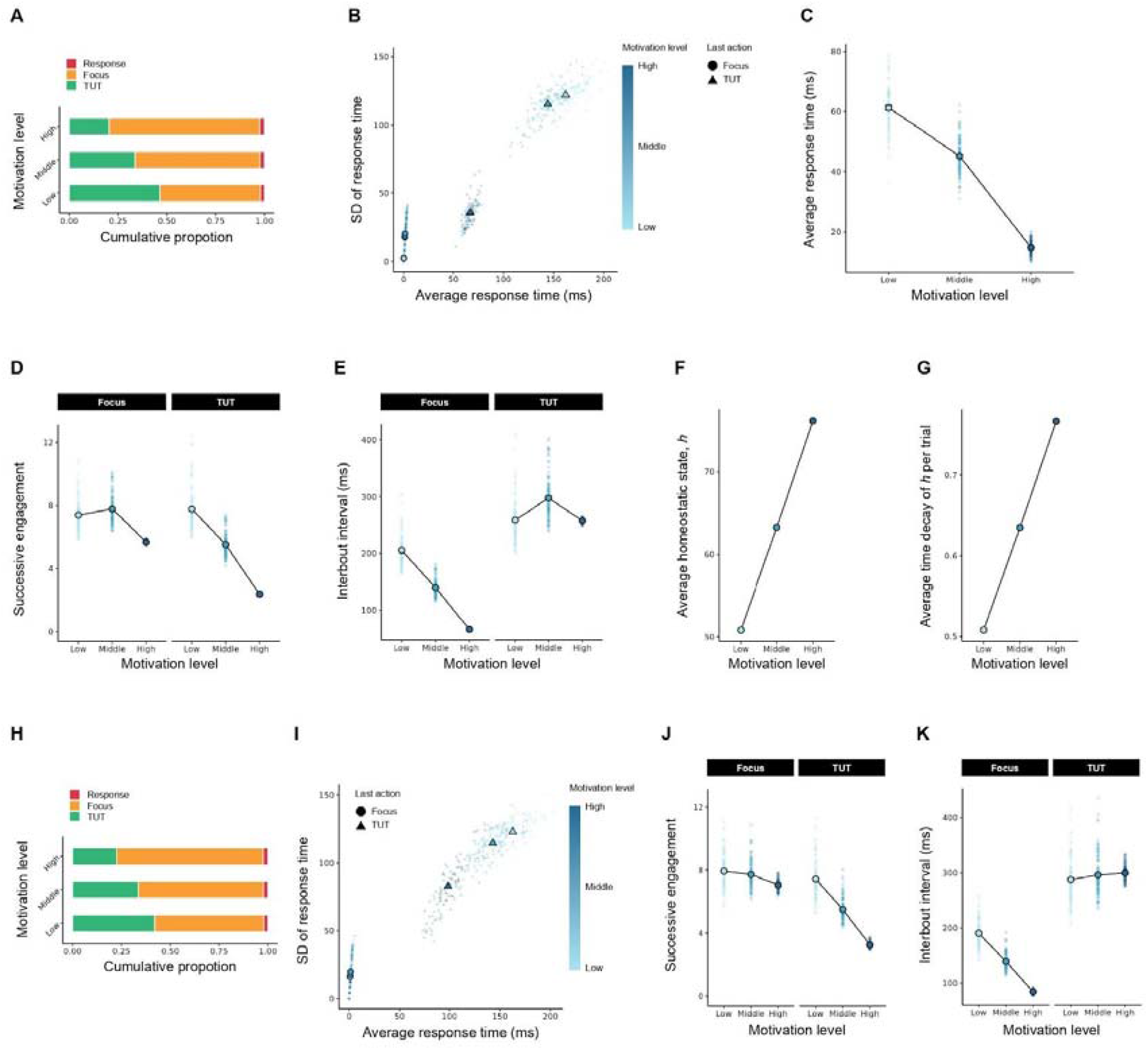
Results of simulation 2-1 and 2-2 We presented the results when the motivation for the task was manipulated in simulations 2-1(A-G) and 2-2(H-K). A, H. Proportion of three actions, “response” (red), “Focus” (orange), and “TUT” (green), when the motivation for the task was manipulated from low (bottom) to high (top). B, I. Average response time and variance upon “stimulus” presentation for each motivation level and agent’s action. The point shape indicates the selected action types upon “stimulus” presentation. As the point color becomes brighter, the motivation for the task decreases. C. Average response time during the task for each motivation level. D, J. Average successive engagement from the action initiation to the end of the action for each motivation level. We counted the number of repetitive choices for each action. E, K. Average inter-action interval duration for each motivation level. F. Average homeostatic state during the task for each motivation level. G. Average time decay during the task for each motivation level. For B–G and I–K, the black-edge points indicate the averaged value and transparent points are individual data averaged across trials.

In Simulation 2-2, the effect of task motivation on the proportion of MW during tasks was also replicated by manipulating the time decay parameters. Similar to the results in Simulation 2-1, as time decay decreased, i.e., decreasing to maintain motivation for the task-related action, the agents engaged more “TUT” (Fig. 3H). Also, the average response time and variance upon “stimulus” presentation increased with the higher motivation level (Fig. 3 I), replicating the relationship between changes in motivation for the task, the proportion of MW, and task performance. The microstructures of behavior, successive engagements, and interbout interval indicated the same trends as the results of simulation 2-1 (Fig 3J, K). Although we manipulated different model parameters, setpoints, and time decay, they influenced agents’ behaviors in the same way, suggesting these two manipulations were functionally equivalent. In summary, we successfully replicated previous experimental findings that the motivation for the task affects the number of MW occurrences and response times and their variance through the manipulation of setpoints and time decay.

In Simulation 3, we simulated the behavior when the setpoint of “TUT” was manipulated and replicated the influence of motivation for TUT on the proportion of MW during tasks. Fig. 4A shows the proportion of action choices for each condition and reveals that HRL agents engaged more “TUT” with increased motivation for the “TUT.” While the response time and variance when agents engaged “TUT” upon “stimulus” presentation increased with higher motivation in simulation 2, we did not observe the effect in simulation 3 (Fig. 4B). However, the average response time during the task prolonged as the setpoint increased (Fig 4C). We conducted further analysis of behavioral microstructures, revealing that setpoint manipulation did not influence successive engagement of actions. In contrast, the interbout interval was decreased as the setpoint for “TUT” became higher (Fig. 4D, E). While the proportion of “TUT” in the task varied across the motivation levels (Fig. 4A), the response time was not prolonged when the agent engaged in “TUT” upon “stimulus” presentation because the motivation level had no effects on the successive engagement (Fig. 4D). Even though the manipulation of the “focus”-setpoint did not change the successive engagement of the “TUT,” the manipulation of the “TUT”-setpoint did change the successive engagement of the focus (Fig. 4D). In Simulation 2-1, the setpoints of “focus” were always set as higher than the TUT, while this simulation included cases where the “TUT” was below or above the “focus”, suggesting that the relative values of the setpoints affect the probability of occurrence for the other behavior. Taken together, we successfully reproduced the effect of MW motivation on behavioral features under the SART.

**Fig. 4.**
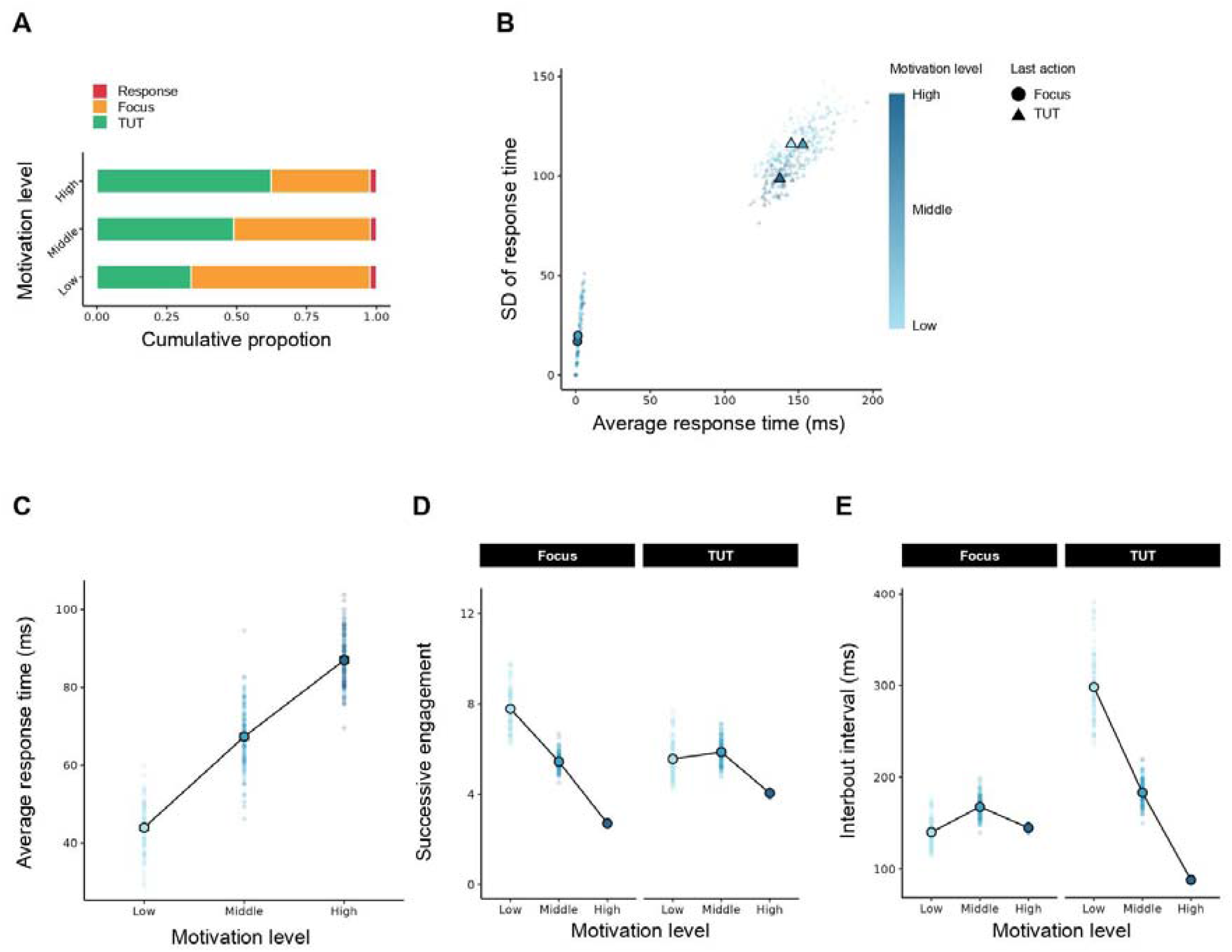
Results of simulation 3 A. Proportion of three actions, “response” (red), “Focus” (orange), and “TUT” (green), when the motivation for the “TUT” was manipulated from low (bottom) to high (top). B. Average response time and variance upon “stimulus” presentation for each motivation level and agent’s action. The shape indicates the selected action types upon “stimulus” presentation. As the point color becomes brighter, the motivation for the task decreases. C. Average response time during the task for each motivation level. D. Average sustained duration when action selection is initiated for each motivation level. E. Average inter-action interval duration for each motivation level. For B–E, the black-edge points indicate the averaged value for each motivation level, and transparent points are individual data averaged across trials.

In Simulation 4, to examine the effect of task difficulty on MW, we simulated the behavior of HRL agents by decreasing the probability of reward for “focus” based on the conditions of Simulation 1 (See Materials and Methods for details). While the proportion of “TUT” decreased as task difficulty increased, conversely, the proportion increased when the task difficulty was extremely high (Fig. 5A). As the task difficulty increased, the response time and variance when the agents engaged in “TUT” upon the “stimulus” presentation, and average response time during the task was decreased with the task difficulty. In contrast, the average response time and its variance were larger than the high level in the extremely high level (Fig. 5B, C). We further analyzed the microstructure of the behavior, and the successive engagement for “focus” did not change monotonically but “TUT” decreased monotonically as the task difficulty increased (Fig. 5D). The interbout interval showed a reversed relationship, that is, “focus” decreased monotonically but “TUT” did not (Fig. 5E). As shown in Fig. A–E, the model displayed non-monotonic changes in behavior for task difficulty and we conducted further analysis on internal variables of the model to clarify underlying mechanisms. When the task was easy, agents could obtain rewards at high probabilities, and it caused saturation in the homeostatic state for “focus” (Fig. 5F). If the homeostatic state for “focus” was saturated, engagement in “TUT” reduced the homeostatic state for “focus” (Fig. 5F) and it produced positive RPEs for “TUT” (Fig. 5 G). In contrast, when the task was difficult, the homeostatic state for “focus” was not saturated. In that case, engagement in “TUT” leads to deviation from the setpoint for “focus” (Fig. 5F), and it caused negative RPEs for “TUT” (Fig. 5G). However, the task was extremely difficult, agents could not obtain a reward at all, and the homeostatic state for “focus” remained constant regardless of agents’ action (Fig. 5F). Thus, the “focus” dimension produced no RPEs for “TUT” (Fig. 5G). The homeostatic state for “focus” converged different asymptotes across task difficulties, and they produced different magnitudes of RPEs from positive to negative. As a result, the task difficulty influenced MW in a non-monotonic way. These results indicate that Simulation 4 succeeded in replicating the effect of task difficulty on MW.

**Fig. 5.**
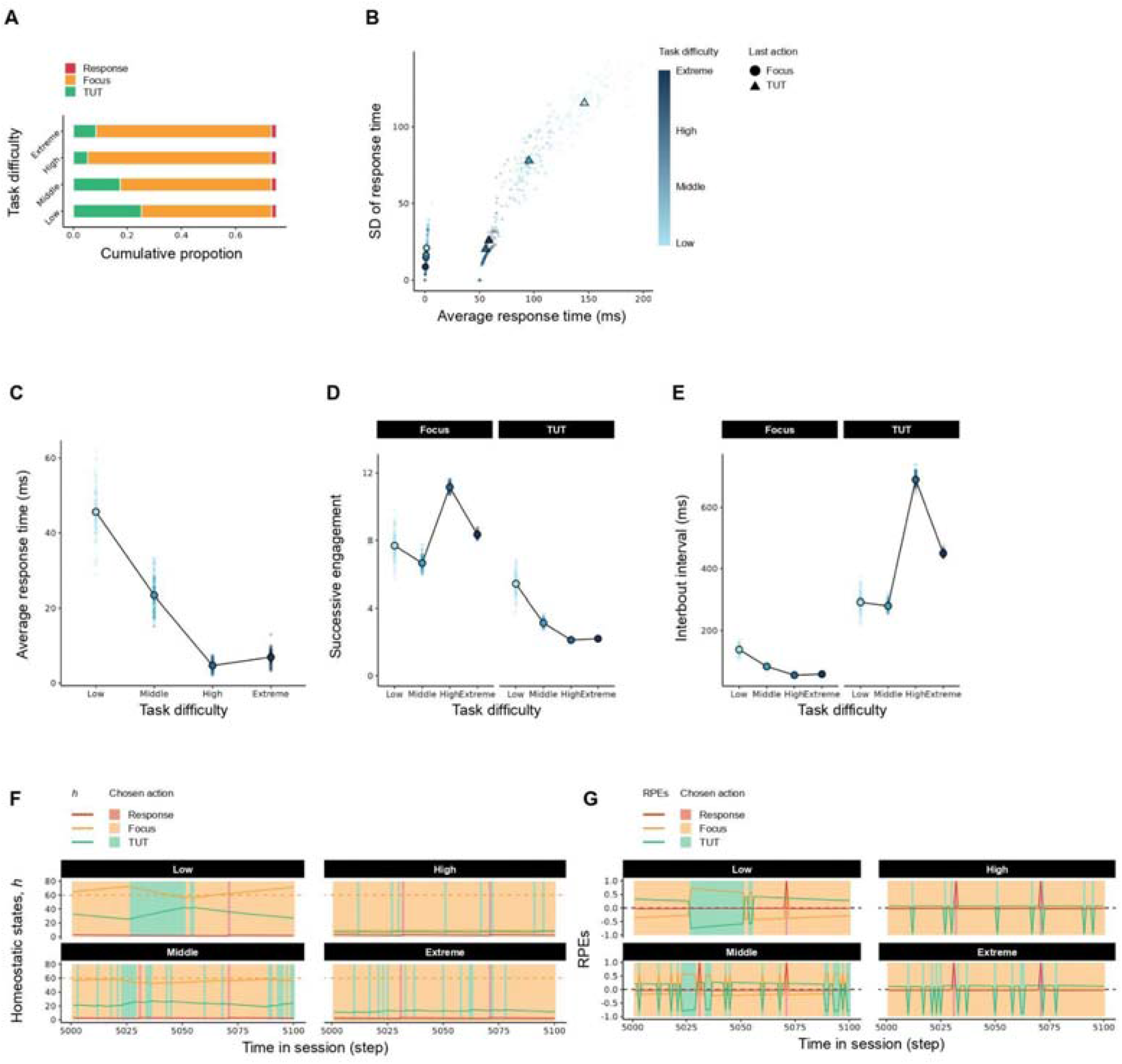
Results of Simulation 4 A. Proportion of three actions, “response” (red), “Focus” (orange), and “TUT” (green), when the motivation for the “TUT” was manipulated from low (bottom) to Extreme (top). B. Average response time and variance upon “stimulus” presentation for each task difficulty and agent’s action. The shape indicates the selected action types upon “stimulus” presentation. As the point color becomes brighter, the task difficulty decreases. C. Average reaction time during the task for each task difficulty. D. Average sustained duration when action selection is initiated for each task difficulty. E. Average inter-action interval duration for each task difficulty. F, G. Time series of homeostatic state and RPEs for each action in a specific agent across some trials. We plotted these values in solid lines and highlighted the interval depending on the actions chosen by agents. For B–E, the black-edge points indicate the averaged value for each motivation level, and transparent points are individual data averaged across trials.

### Interim Discussion

In Simulation 2∼4, we qualitatively replicated the experimental results presented in MW studies and showed differences in HRL behavior following the manipulation of parameters. In Simulation 2 and 3, we manipulated the motivation for the task and TUT by manipulating the setpoint of each action and the parameters governing the time decay in the homeostatic state, replicating the changes in response time, variance, and the proportion of MW during task reported in previous studies. In Simulation 4, we changed the task difficulty by manipulating the reward probability for “focus.” The results indicated that the proportion of MW decreased with increasing task difficulty but increased at extremely high difficulty levels. These results validate the assumption that HRL provides a comprehensive explanation of the main experimental results related to MW occurrence and becomes an underlying mechanism of MW.

In addition to the experimental results of MW validated through the simulations, the traits of MW observed in the HRL agents are consistent with many findings in the previous studies. At the onset of MW in HRL, the reward for focusing on the main task is negative, and the TUT is driven to resolve this saturated homeostatic state. This functional aspect of MW is also consistent with the hypothesis that MW has a distracting effect that may alleviate the current boredom (Mooneyham and Schooler, 2013). Temporal patterns in the proportion of MW within a session showed a low occurrence rate initially, which increased as the task continued, and similar results have been reported in experimental settings (Brosowsky et al., 2020; Zanesco et al., 2024). The decrease in the proportion of MW among older adults can be explained by fewer cues related to current concerns in the experimental setting (McVay et al., 2013), consistent with the findings when we manipulated motivation for TUT in the present study. Although not explicitly addressed in this study, HRL can also explain various aspects of MW.

Characteristics of thought processing, such as intentionality or automaticity, are considered important in MW research (Christoff et al., 2016). Although the model presented in this study does not explicitly address these aspects due to the agent’s inherent lack of subjectivity, it provides some valuable insights. Two types of MWs emerged in this model: those intended to alleviate excessive engagement to the main task (Simulation 1) and those intended to actively engage in MWs driven to fill in the gap between the current homeostatic state and setpoint (Simulation 3). Intentional MW tends to relate the contents to the future (Seli et al., 2017). Given that we assumed the active MW was driven by current concerns in Simulation 3, this result suggests that active engagement in MW may be associated with intentional thought. In addition, unintentional MWs occurred more frequently over task time than intentional MWs (Martínez-Pérez et al., 2021). These results indicated that the unintentional MWs may be consistent with the characteristics of MWs, which is intended to alleviate excessive engagement in the main task and support the correspondence between intentionality and the MW types of the HRL model.

The microstructure analysis revealed parameter manipulations that influence which aspect of MW and it enabled us to compare the results of our simulations and previous findings in detail. The previous study has shown that higher motivation for the task decreases MW by mediating the effort to return attention to the task more quickly (He et al., 2023), suggesting that the duration of each TUT is shortened. When we manipulated motivation for the task in simulation 2, the proportion of MW decreased as the task motivation increased and it was mediated by shorter successive engagement in “TUT” (Fig 2E and J), suggesting that HRL could capture the underlying mechanism behind thought transition. This is not limited to MW, but animal behavior showed a similar relationship between motivation and behavioral microstructure. Instrumental behaviors of animals show bout-and-pause patterns that are characterized by bursts of instrumental responses and longer pauses separating each burst (Gilbert, 1958; Shull et al., 2001). It is reported that motivation, such as hunger, controlled interbout intervals, and bout length (Podlesnik et al., 2006; Shull, 2004; Shull et al., 2001). These results suggest that MWs have a similar microstructure to those observed in observable animal behavior, supporting the idea that thought phenomena share a common principle with the general behavior of organisms.

### General Discussion

The present study aimed to comprehensively explain the previous results related to MW occurrence using HRL through the simulations and provide evidence that action selection through homeostatic control is behind the dynamics of thoughts. We imposed the SART on HRL agents and examined response time, behavioral microstructures, and the internal variables as a function of parameter manipulations. The results showed that MW in HRL agents occurred when the homeostatic state for the task-focused was saturated, and the relative action value function of TUT increased (Fig. 2). Furthermore, we successfully replicated the effects of motivation for the task and MW, and task difficulty on the proportion of MW reported in previous studies by manipulating the relevant parameters (Figs 3-5). In summary, HRL could provide a comprehensive explanation for MW, suggesting that human thought transitions follow the homeostatic control of action selection, that is, choices between the task and TUT and independent drives dynamics behind them.

One of the concerns with employing HRL to provide an account for MW is that the model handles an observable action (“response”) and unobservable ones (“focus” and “TUT”) in the same line. However, this formulation is validated by the following three points: 1) Many MW studies recruited probe-caught methods, which ask participants to choose whether they have focused on the task or MW randomly during a task (Weinstein, 2018). This method is similar to the present study, where MW is treated along the same lines as task engagement as participants’ thought options. 2) The fact that MW prolonged the response time in our simulations and empirical studies, indicates that the TUT competes with observable response or task-focused states even in the behavioral output. 3) The hypothesis that MW uses the same cognitive resources as task engagement has long been proposed (Smallwood and Schooler, 2006) and is supported by many psychological and neuroscientific findings (Christoff et al., 2009; Randall et al., 2019). These findings and assumptions indicate that previous studies implicitly or explicitly treated internal thoughts on the same level as other observable behaviors, especially in assessing whether the agents were engaged in the task.

Another concern is to set a homeostatic state or setpoint for task-related actions and TUT. In the present study, the homeostatic state of behavior does not assume a corresponding physiological state, as with conventional HRL. However, several perspectives in prior research support the idea of treating the amount of engagement in a particular behavior as the homeostatic state without corresponding to the physiological state and that action choices minimize deviations of this state from setpoint. The motivation hypothesis, which explains the occurrence of MW, hypothesized that psychological tasks are simple and task engagement restricts other behaviors, resulting in lost opportunities and an increase in the action value of alternative behaviors, the accumulation of which leads to attentional shift (Brosowsky et al., 2020). This perspective implicitly assumes that task engagement depends on the comparisons between the value of TUT and task engagement calculated from the amount of engagement and the threshold. Thought transitions in the neuroscientific perspective are explained by the fact that continued activity in the locus coeruleus, a brain region that contributes to sustained attention, leads to hyperactivity and a release of the task-focused state beyond the optimal activity point (Mittner et al., 2016). Although the process after the release of focus is unknown, a threshold of activity is assumed for the locus coeruleus, which would lead to the termination of a specific thought. Extending homeostasis beyond physiological states, such that engagement in a particular behavior or thought has some threshold for the amount of engagement, does not deviate from the assumptions and findings of previous studies. Indeed, some studies postulate homeostasis in social interactions and habituation (Eisenstein and Eisenstein, 2006; Lee et al., 2021).

The explanation of MW using HRL also includes a perspective on interactions between brain regions among the thought transitions frequently discussed in the context of MW research. Three brain regions are thought to be related to task engagement and MW: 1) the central executive network (CEN), which is active during task engagement, 2) the default mode network (DMN), which is associated with internal thought, and 3) the salience network (SN) switching the transition between the two networks (Christoff et al., 2016; Menon and Uddin, 2010; Molnar-Szakacs and Uddin, 2022). The transition from MW to task-focused state is explained by the SN detecting salient stimulus in the internal and external environment and switching to CEN. However, the transition from the focused state to MW, i.e., switching when the DMN is activated, has yet to be fully explained (Schimmelpfennig et al., 2023; Seeley, 2019; Sridharan et al., 2008). The insular cortex, which constitutes the SN, is known to be involved with emotional and physical responses (Schimmelpfennig et al., 2023). This region is also activated immediately before action selection changes, especially in environments with little change (Munn et al., 2021; Yawata et al., 2023). Our simulations revealed that the transitions from task engagement to TUT and from TUT to task engagement are driven by saturation of the currently engaged action (Fig. 2). Assuming that the SN tracks the internal state of each dimension of action, HRL can provide a comprehensive account of bidirectional thought transitions, such as the occurrence of MW (the activation of DMN) and the engagement in the task (the activation of CEN) in the form of the detection of agents’ unpleasant emotions and their associated physical responses by the SN. Furthermore, HRL has the potential to include the case that salient stimulus detections occur suddenly in the internal and external environment, which conventional brain network models have assumed. In this study, as the task time progressed, the internal states of both “focus” and “TUT” deviated from the setpoint, and the action value function converged to a negative value (Fig 2). When a salient novel stimulus emerges, the action value function of that stimulus is relatively high compared to other alternatives. Thus, when any internal state dimension is saturated, it is possible to detect and process a salient stimulus. Studies in humans have shown that the degree of task engagement, which is manipulated by fatigue and motivation, affects the degree of suppression of task-irrelevant stimuli, supporting our hypothesis (Buetti and Lleras, 2016; Faber et al., 2012). The behavior of HRL is consistent with the action-level shifts between MW and task engagement and the interactions among the relevant brain networks.

We showed that HRL, a model for associative learning of animals, could reproduce MW, suggesting that thoughts unique to humans could be explained from the view of animal behavior. Animals must satisfy several requirements, such as hunger, thirst, danger, and mating in natural environments. The facts lead us to assume that there can be a different principle from maximizing a single reward, as expected in a laboratory setting. Evidence for this has long been reported in animal behavior research, such as engaging in diverse behaviors not related to the experimental task (Breland and Breland, 1961; Falk, 1966; Gentry, 1968; Levitsky and Collier, 1968; Skinner, 1948), and changes in the relative reward effects of behavioral opportunities, such as a less preferred behavior reinforce a more preferred behavior under specific environmental settings (Allison and Timberlake, 1974), have been reported, and it has been theoretically suggested that task-unrelated behaviors influence the behavior of task-related behaviors Theories suggest that they may influence task-related behavioral behavior (Baum, 2012; Guthrie, 1930; Herrnstein, 1970; Killeen and Fetterman, 1988; Staddon, 1979; Timberlake and Allison, 1974; Yamada and Toda, 2023). In addition, we showed that behavioral changes caused by motivational operation to HRL were similar to those of animal operant behaviors (Podlesnik et al., 2006; Shull, 2004; Shull et al., 2001). Taken together, there can be a similar principle behind human thoughts and transitions between them to those of animal behaviors. Boredom-like behavior in which animals and humans tend to accept aversive stimulus, such as electric shocks or air puffs, in a poor environment, and its neural substrates support our idea (Wilson et al., 2014; Yawata et al., 2023). When the environment is poor, agent homeostatic states for available actions should satiate, and aversive stimulus should become a reward by reducing satiation. In such a situation, the insular cortex, a part of SN, was involved in the boredom-like behavior, and mesolimbic dopaminergic neurons were excited around the aversive stimulus presentations (Yawata et al., 2023). We showed that “TUT” reduced the saturation of the homeostatic state for “focus,” and it works as a reward for “TUT” (Fig. 2K, L). The process that drives “TUT” is similar to the mechanism of boredom-like behavior of mice. This is not limited to conceptual similarity but neural evidence that the insular cortex is involved in boredom-like behavior, and SN supports our idea. Thus, our results suggest that animal behaviors and the dynamics of thought, a phenomenon unique to humans, may have a common basis in homeostatic behavior control.

In daily life, we get lost in thoughts of various matters, such as issues or problems we face, a future of ourselves, and past mistakes. What drives these thoughts to transition from one thing to another? Here, we provide a clear answer to the question. The difference between the setpoint and the amount of engagement for options decides the action to be engaged at the next step. Despite the simplicity, we successfully replicated the previous findings in the MW research through our simulations and provided a comprehensive explanation for MW. Although not addressed in this study, it has high applicability to spontaneous thought transition phenomena in general since it can be extended not only to bidirectional transitions between task and MW but also to sudden events as described above and to more segmented thoughts of TUT, i.e., to assume each dimension for the future and the past. Furthermore, we revealed the similarity between the process of MW occurrence and animal behavior by analyzing the internal dynamics of the agents. Taken together, we could explain human thought transition by extending a model of animal learning and behavior based on homeostatic control. This suggests that human thought shares a more fundamental principle with animal behavior than we had usually assumed.

## Methods

### Sustained attention to response task in the simulation

Many MW studies used SART as the main task, in which fixation and numbers (e.g., 1∼9) are presented alternately, and participants are asked to press a key as quickly as possible when the specific number (e.g., 5) appears (Fig. 2A). Therefore, the participants need to attend and respond to the numbers presented at all times, which requires sustained attentional allocation to the task. At random time points during the task, participants are asked about their attentional state, such as whether they were focused on the task or MW. By measuring the response time, brain activity, and physiological indices before the MW reports, the previous research revealed the traits of MW. Many studies have pointed out the delay of response time and the increase in the variance during MW, we used these indices as the measure of MW in the present study. We used a simplified version of this experiment because the main focus is describing the relationship between action choice and the internal states underlying it with manipulating parameters. Agents were presented with the “fixation” or “stimulus” at each time step and required to respond as quickly as possible when the task state changed to “stimulus.” In this situation, the agents focused on the task engaged TUT or responded to the “stimulus” at each time point (Fig. 2B, C).

The inter-stimulus interval (ISI), the presentation time of the “fixation,” was 1500 ms, the “stimulus” presentation time was 500 ms, and each timestep was 50 ms, resulting in 40 steps per trial. In all simulations, 200 trials were conducted, i.e., 2000 ms x 200 trials = 400 seconds (about 6.5 minutes). We ran 100 simulations with the same setting for each condition in all simulation sections, assuming we collected data from 100 participants. Since ISI is 30 steps (1500/50 ms), the agents could respond when the time step is between 31 to 40 steps. When we presented the results of simulations, time steps were changed into actual time in seconds.

### Homeostatic reinforcement learning

HRL aims to minimize the deviation of the internal homeostatic state from the setpoint, assuming homeostasis is maintained throughout the reinforcement learning. The agent’s physiological state, or homeostatic state, is formulated *H*_*i,t*_ = (*h*_*l,t*_ *h*_2,*t*_ … *h*_*N,t*_) and defined as a multidimensional space consisting of time-varying body temperature, blood glucose level, blood pressure, etc. The *i*-th internal state at a time point *t* is denoted by_i,*t*_. Each homeostatic state decays exponentially through time according to the following;

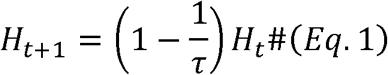

*τ* is a time constant that defines the decay speed in Eq. 1. In this homeostatic space, the drive function *D*(*H*_*t*_) was defined as the distance of the homeostatic state from the setpoint 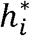 as follow;

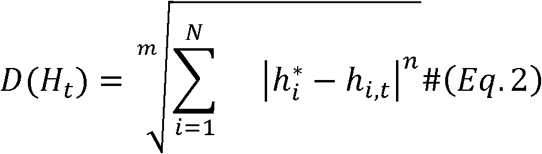

In Eq. 2, *m* and *n* are free parameters that induce nonlinear effects on the mapping between homeostatic deviations and their motivational consequences. Rewards are defined as a reduction in drives when agents get observation from environments, implying that changes in homeostatic space work as rewards. The reward is defined as following;

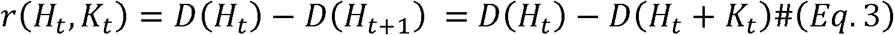

To maximize cumulative reward over time, agents must learn state–action mapping. The Q-learning model was employed to learn action value functions. In this model, the values of action *Q*_*t*_ (*a*) is updated based on the reward prediction error.

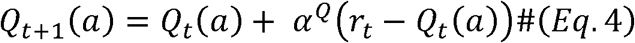

In Eq. 4, *α* is the learning rate for action values and determines how significantly the prediction error is evaluated. The choice are decided depends on the probability calculated from the soft-max function.

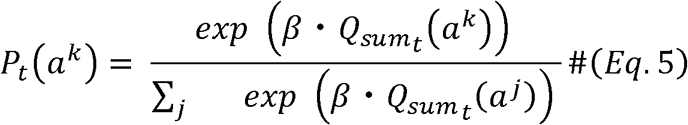

*P*_*t*_ (*a*^*k*^) is the probability of an action to be selected at time *t*, and *β*, is the inverse temperature, which controls the randomness of choice. The overarching architecture of HRL is outlined above, with several processes added to the conventional HRL model to implement the SART on agents (Fig. 2D).

### Homeostatic reinforcement learning for sustained attention to response task

Here, we describe how we apply HRL to the SART. SART involves two different environment states, “fixation” and “focus,” and available actions differ in the environment states. Agents can choose three actions, “response,” “focus,” and “TUT.” “Response” is the response to the stimulus, “focus” is the agent engaged in the task but not responding to the stimulus, and “TUT” is a task-unrelated thought. “Response” is only available when the environment state is “stimulus,” but others are available in both states. To impose constraints on agents’ choice depending on the environment state, we set different transition probability matrices for each state and previous action choice and weighted the action values calculated by Eq. 4 (Fig. 2D). Also, the agents were required to focus on the task constantly, so we modeled using shared action values across states of the environment to ensure that the action value function does not change depending on whether the stimulus is presented.

We also assumed an additional process to the HRL, that is the persistence of action. In general, thoughts and actions are not considered to be in immediate and frequent transition but rather to be maintained through time (Bastian and Sackur, 2013; Shinagawa et al., 2023; Zanesco et al., 2020). We incorporate the persistence to the HRL by having agents to choose action based on not only action values but previous actions. First, we introduce an action trace, which represents how many times each action has been selected in the past timesteps. The perseverance, denoted as 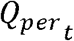, is calculated by the following equation;

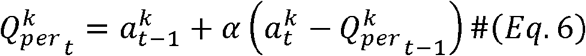

In Eq. 6, *α* is a free parameter that defines how much weighting the immediate before the response. The Action, *α*_*t*_, is represented by a binary, if the action *k* is chosen in a timestep *t*, then *a*^*k*^ is 1 otherwise is 0. If agents choose the action *k* repeatedly, 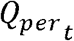 is increased, otherwise it decreases. Finally, agents choose their action based on the q-values (Eq. 4) and persistence (Eq. 6) by calculating the weighted sum of them as follows;

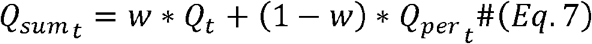

### Parameter settings for each simulation

#### Simulation1

We showed the specific parameter settings used in each simulation. Simulation 1 was conducted to reveal that MW occurrence depends on two independent processes: 1) a choice between task focus and TUT, and 2) independent dynamics of the drive behind them. To achieve this goal, we simulated and analyzed the behavior under SART of three models: a full model incorporating both processes, a no-TUT model, and a vanilla Q-learning model (VQ). In the full model where an action choice includes “TUT,” the transition probabilities for “focus” and “TUT” were set to be the same during “fixation,” and which one is chosen depends on the action value (Table. 1). In a no-TUT model, the overall structure was the same as the full model but controlled using transition probabilities to restrict the choice of “TUT.” We set the transition probability to select “TUT” as 0 from the task-related actions (Table. 2). When the agents selected a “response,” we applied the same transition probability as the “fixation” to avoid consecutive responses to prevent agents from choosing “response” many times in one trial. We also examined the behavior of a VQ in SART to show the independent dynamics of the drive behind “focus” and “TUT,” which is a necessary process for MW occurrence. In the VQ model, there was no change in the homeostatic states so rewards were clipped at constant through time. We set the observed reward from the environment to 1. for all models and simulations.

**Table. 1.** Transition probabilities when task-unrelated thoughts are included in the action options.

| Environment state |  | Response | Focus | TUT |
| --- | --- | --- | --- | --- |
| Fixation | Response | 0. | .5 | .5 |
|  | Focus | 0. | .5 | .5 |
|  | TUT | 0. | .5 | .5 |
| Stimulus | Response | 0. | .5 | .5 |
|  | Focus | .5 | .25 | .25 |
|  | TUT | 0. | .5 | .5 |

**Table. 2.** Transition probabilities in environments where task-unrelated thought cannot be selected.

| Environment state |  | Response | Focus | TUT |
| --- | --- | --- | --- | --- |
| Fixation | Response | 0. | 1. | 0. |
|  | Focus | 0. | 1. | 0. |
| Stimulus | Response | 0. | 1. | 0. |
|  | Focus | .5 | .5 | 0. |

The other parameter settings of the HRL agent used in Simulation 1 are shown in Tables 3 and 4. First, parameters such as learning rate, m, and n were determined by referring to previous studies, while the inverse temperature, the learning rate, and the weight parameter (w) for perseverance were determined based on the simulation results (Table 3). In all simulations, the initial values of the homeostatic state and Q-value were set to 0, so the first action was randomly chosen. Furthermore, Table 4 shows the settings of the setpoints and time decay for each action choice. The homeostatic state of “TUT” is equal to the setpoint from the onset of the task, meaning that the drive for “TUT” always becomes negative during the task.

**Table. 3.** Parameter settings related to learning of HRL agents.

| | $\alpha$ | $\beta$ | $m$ | $n$ | $w$ |
| --- | --- | --- | --- | --- | --- |
| values | .05 | 10 | 3 | 4 | .95 |

**Table. 4.** Other parameter settings in simulation 1.

|  | Response | Focus | TUT |
| --- | --- | --- | --- |
| tau | 1/100 | 1/100 | 1/100 |
| setpoint | 30 | 30 | 0 |

#### Simulation2

To replicate the finding that motivation for the task reduces the proportion of MW, we manipulated the setpoint (Table. 5) and the speed of spontaneous attenuation in the homeostatic state (Table 6) for the task-related actions. Since higher setpoints lead to a larger deviation between the homeostatic state and setpoints, this parameter could directly control the motivation. Time decay specifies the drive’s recovery speed and seemed to be motivation. The setpoints and time decay used in Simulation 1 were set as the reference (Mid), and we manipulated this value to lower (Low) or higher values (High). The other parameters are the same as in the full model used in Simulation 1.

**Table. 5.**
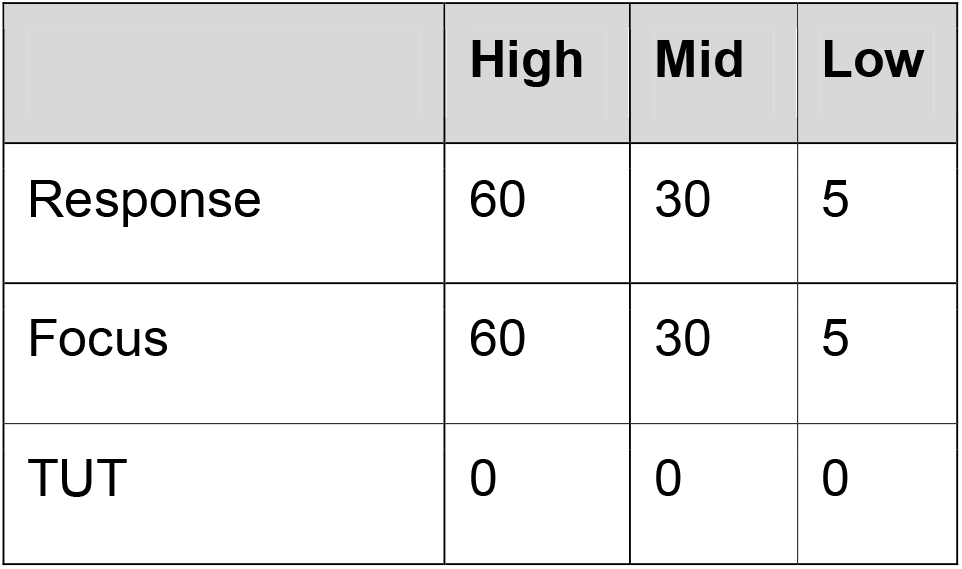
Setpoints for task related actions in each condition in Simulation2-1.

|  | High | Mid | Low |
| --- | --- | --- | --- |
| Response | 60 | 30 | 5 |
| Focus | 60 | 30 | 5 |
| TUT | 0 | 0 | 0 |

**Table. 6.** Time decay for task related in each condition in Simulation2-2.

|  | High | Mid | Low |
| --- | --- | --- | --- |
| tau | 1/10 | 1/100 | 1/1000 |

#### Simulation3

To replicate the finding that highly concerned events drive MW, we manipulated the setpoint to the “TUT” (Table. 7). As mentioned above, the setpoints directly control the motivation for the action. We increased only the setpoint for “TUT” from that used in the full model in Simulation 1, and the other parameters were the same.

**Table. 7.** Setpoints for task-unrelated thought in each condition in Simulation3.

|  | Response | Focus | TUT |
| --- | --- | --- | --- |
| low | 30 | 30 | 0 |
| mid | 30 | 30 | 30 |
| high | 30 | 30 | 60 |

#### Simulation4

To replicate the relationship between the proportion of MW and the task difficulty, we manipulated the change probability of homeostatic state depending on the action choice (Table 8). The agent does not fulfill the task as expected by manipulating the probability of homeostatic state updates. We tested the effect of task difficulty by setting the probability used in the full model of Simulation 1 to the reference condition (Low) and reducing the probability from there. In addition, to reproduce a situation in which the task was extremely difficult, we used a condition in which the probability of homeostatic state change was set to 0 in which the agents did not feel fulfilled no matter how much they engaged in the task. The other parameters are the same as in the full model of Simulation 1.

**Table. 8.** Setting the probability of homeostatic state change for the action choice for each condition of Simulation 4.

|  | Extreme | High | Mid | Low |
| --- | --- | --- | --- | --- |
| Response | 1. | 1. | 1. | 1. |
| Focus | 0. | .1 | .75 | 1. |
| TUT | 1. | 1. | 1. | 1. |

## Code availability

The codes generating and analysing data were available via GitHub at https://github.com/shina-k/HRL-MW.git.

## Data availability

All relevant data are within the paper (Figs. 2–5) and the data and figures were generated using author’s scripts (See Code availability).

## Acknowledgements

This research was supported by JSPS KAKENHI 24K16869 (KY) and 24KJ0069 (KY).

## Author information

These authors contributed equally: Kazushi Shinagawa, Kota Yamada.

### Contributions

K.S., and K.Y. conceived the idea for the paper, analyzed data and wrote and edited the paper.

## Reference

Allison J, Timberlake W. 1974. Instrumental and contingent saccharin licking in rats: Response deprivation and reinforcement. Learn Motiv 5:231–247.

Baird B, Smallwood J, Schooler JW. 2011. Back to the future: autobiographical planning and the functionality of mind-wandering. Conscious Cogn 20:1604–1611.

Barrington M, Miller L. 2023. Mind wandering and task difficulty: The determinants of working memory, intentionality, motivation, and subjective difficulty. Psychology of Consciousness: Theory, Research, and Practice. doi:10.1037/cns0000356

Bastian M, Sackur J. 2013. Mind wandering at the fingertips: automatic parsing of subjective states based on response time variability. Front Psychol 4:573.

Baum WM. 2012. Rethinking reinforcement: allocation, induction, and contingency. J Exp Anal Behav 97:101–124.

Breland K, Breland M. 1961. The misbehavior of organisms. Am Psychol 16:681–684.

Brosowsky NP, DeGutis J, Esterman M. 2020. Mind wandering, motivation, and task performance over time: Evidence that motivation insulates people from the negative effects of mind wandering. Psychology of.

Buetti S, Lleras A. 2016. Distractibility is a function of engagement, not task difficulty: Evidence from a new oculomotor capture paradigm. J Exp Psychol Gen 145:1382–1405.

Christoff K, Gordon AM, Smallwood J, Smith R, Schooler JW. 2009. Experience sampling during fMRI reveals default network and executive system contributions to mind wandering. Proc Natl Acad Sci U S A 106:8719–8724.

Christoff K, Irving ZC, Fox KCR, Spreng RN, Andrews-Hanna JR. 2016. Mind-wandering as spontaneous thought: a dynamic framework. Nat Rev Neurosci 17:718–731.

Eisenberger R, Karpman M, Trattner J. 1967. What is the necessary and sufficient condition for reinforcement in the contingency situation? J Exp Psychol 74:342–350.

Eisenstein EM, Eisenstein D. 2006. A behavioral homeostasis theory of habituation and sensitization: II. Further developments and predictions. Rev Neurosci 17:533–557.

Faber LG, Maurits NM, Lorist MM. 2012. Mental fatigue affects visual selective attention. PLoS One 7:e48073.

Falk JL. 1966. Schedule-induced polydipsia as a function of fixed interval length. J Exp Anal Behav 9:37–39.

Gentry WD. 1968. Fixed-ratio schedule-induced aggression. J Exp Anal Behav 11:813–817.

Gilbert TF. 1958. Fundamental dimensional properties of the operant. Psychol Rev 65:272–282.

Guthrie ER. 1930. Conditioning as a principle of learning. Psychol Rev 37:412.

He H, Chen Y, Li T, Li H, Zhang X. 2023. The role of focus back effort in the relationships among motivation, interest, and mind wandering: an individual difference perspective. Cogn Res Princ Implic 8:43.

Henríquez RA, Chica AB, Billeke P, Bartolomeo P. 2016. Fluctuating Minds: Spontaneous Psychophysical Variability during Mind-Wandering. PLoS One 11:e0147174.

Herrnstein RJ. 1970. On the law of effect. J Exp Anal Behav 13:243–266.

Irrmischer M, van der Wal CN, Mansvelder HD, Linkenkaer-Hansen K. 2018. Negative mood and mind wandering increase long-range temporal correlations in attention fluctuations. PLoS One 13:e0196907.

Juechems K, Summerfield C. 2019. Where Does Value Come From? Trends Cogn Sci 23:836–850.

Kachanoff R, Leveille R, McLelland JP, Wayner MJ. 1973. Schedule induced behavior in humans. Physiol Behav 11:395–398.

Kahmann R, Ozuer Y, Zedelius CM, Bijleveld E. 2022. Mind wandering increases linearly with text difficulty. Psychol Res 86:284–293.

Keramati M, Gutkin B. 2014. Homeostatic reinforcement learning for integrating reward collection and physiological stability. Elife 3. doi:10.7554/eLife.04811

Killeen PR, Fetterman JG. 1988. A behavioral theory of timing. Psychol Rev 95:274–295.

Killingsworth MA, Gilbert DT. 2010. A wandering mind is an unhappy mind. Science 330:932.

Lee CR, Chen A, Tye KM. 2021. The neural circuitry of social homeostasis: Consequences of acute versus chronic social isolation. Cell 184:1500–1516.

Leszczynski M, Chaieb L, Reber TP, Derner M, Axmacher N, Fell J. 2017. Mind wandering simultaneously prolongs reactions and promotes creative incubation. Sci Rep 7:10197.

Levitsky D, Collier G. 1968. Schedule-induced wheel running. Physiol Behav 3:571–573.

Makovac E, Fagioli S, Watson DR, Meeten F, Smallwood J, Critchley HD, Ottaviani C. 2019. Response time as a proxy of ongoing mental state: A combined fMRI and pupillometry study in Generalized Anxiety Disorder. Neuroimage 191:380–391.

Martínez-Pérez V, Baños D, Andreu A, Tortajada M, Palmero LB, Campoy G, Fuentes LJ. 2021. Propensity to intentional and unintentional mind-wandering differs in arousal and executive vigilance tasks. PLoS One 16:e0258734.

McVay JC, Meier ME, Touron DR, Kane MJ. 2013. Aging ebbs the flow of thought: adult age differences in mind wandering, executive control, and self-evaluation. Acta Psychol 142:136–147.

Menon V, Uddin LQ. 2010. Saliency, switching, attention and control: a network model of insula function. Brain Struct Funct 214:655–667.

Mittner M, Hawkins GE, Boekel W, Forstmann BU. 2016. A Neural Model of Mind Wandering. Trends Cogn Sci 20:570–578.

Molnar-Szakacs I, Uddin LQ. 2022. Anterior insula as a gatekeeper of executive control. Neurosci Biobehav Rev 139:104736.

Mooneyham BW, Schooler JW. 2013. The costs and benefits of mind-wandering: a review. Can J Exp Psychol 67:11–18.

Munn BR, Müller EJ, Wainstein G, Shine JM. 2021. The ascending arousal system shapes neural dynamics to mediate awareness of cognitive states. Nat Commun 12:1–9.

Podlesnik CA, Jimenez-Gomez C, Ward RD, Shahan TA. 2006. Resistance to change of responding maintained by unsignaled delays to reinforcement: a response-bout analysis. J Exp Anal Behav 85:329–347.

Poerio GL, Totterdell P, Miles E. 2013. Mind-wandering and negative mood: does one thing really lead to another? Conscious Cogn 22:1412–1421.

Randall JG, Beier ME, Villado AJ. 2019. Multiple routes to mind wandering: Predicting mind wandering with resource theories. Conscious Cogn 67:26–43.

Schimmelpfennig J, Topczewski J, Zajkowski W, Jankowiak-Siuda K. 2023. The role of the salience network in cognitive and affective deficits. Front Hum Neurosci 17:1133367.

Seeley WW. 2019. The Salience Network: A Neural System for Perceiving and Responding to Homeostatic Demands. J Neurosci 39:9878–9882.

Seli P, Cheyne JA, Smilek D. 2013. Wandering minds and wavering rhythms: linking mind wandering and behavioral variability. J Exp Psychol Hum Percept Perform 39:1–5.

Seli P, Konishi M, Risko EF, Smilek D. 2018. The role of task difficulty in theoretical accounts of mind wandering. Conscious Cogn 65:255–262.

Seli P, Ralph BCW, Konishi M, Smilek D, Schacter DL. 2017. What did you have in mind? Examining the content of intentional and unintentional types of mind wandering. Conscious Cogn 51:149–156.

Shinagawa K, Itagaki Y, Umeda S. 2023. Coexistence of thought types as an attentional state during a sustained attention task. Sci Rep 13:1581.

Shull RL. 2004. Bouts of responding on variable□ interval schedules: Effects of deprivation level. J Exp Anal Behav 81:155–167.

Shull RL, Gaynor ST, Grimes JA. 2001. Response rate viewed as engagement bouts: effects of relative reinforcement and schedule type. J Exp Anal Behav 75:247–274.

Skinner B. 1948. Superstition in the pigeon. J Exp Psychol 38:168–172.

Smallwood J, Beach E, Schooler JW, Handy TC. 2008. Going AWOL in the brain: mind wandering reduces cortical analysis of external events. J Cogn Neurosci 20:458–469.

Smallwood J, Schooler JW. 2015. The science of mind wandering: empirically navigating the stream of consciousness. Annu Rev Psychol 66:487–518.

Smallwood J, Schooler JW. 2006. The restless mind. Psychol Bull 132:946–958.

Sridharan D, Levitin DJ, Menon V. 2008. A critical role for the right fronto-insular cortex in switching between central-executive and default-mode networks. Proc Natl Acad Sci U S A 105:12569–12574.

Staddon JER. 1979. Operant behavior as adaptation to constraint. J Exp Psychol Gen 108:48–67.

Thomson DR, Besner D, Smilek D. 2013. In pursuit of off-task thought: mind wandering-performance trade-offs while reading aloud and color naming. Front Psychol 4:360.

Timberlake W, Allison J. 1974. Response deprivation: An empirical approach to instrumental performance. Psychol Rev 81:146–164.

Uchida Y, Hikida T, Yamashita Y. 2022. Computational Mechanisms of Osmoregulation: A Reinforcement Learning Model for Sodium Appetite. Front Neurosci 16:857009.

Weinstein Y. 2018. Mind-wandering, how do I measure thee with probes? Let me count the ways. Behav Res Methods 50:642–661.

Wilson TD, Reinhard DA, Westgate EC, Gilbert DT, Ellerbeck N, Hahn C, Brown CL, Shaked A. 2014. Just think: The challenges of the disengaged mind. Science 345:75–77.

Xu J, Metcalfe J. 2016. Studying in the region of proximal learning reduces mind wandering. Mem Cognit 44:681–695.

Yamada K, Toda K. 2023. Habit formation viewed as structural change in the behavioral network. Commun Biol 6:303.

Yawata Y, Shikano Y, Ogasawara J, Makino K, Kashima T, Ihara K, Yoshimoto A, Morikawa S, Yagishita S, Tanaka KF, Ikegaya Y. 2023. Mesolimbic dopamine release precedes actively sought aversive stimuli in mice. Nat Commun 14:2433.

Zanesco AP, Denkova E, Jha AP. 2024. Mind-wandering increases in frequency over time during task performance: An individual-participant meta-analytic review. Psychol Bull. doi:10.1037/bul0000424

Zanesco AP, Denkova E, Witkin JE, Jha AP. 2020. Experience sampling of the degree of mind wandering distinguishes hidden attentional states. Cognition 205:104380.

